# Ultrasound Echogenicity is Associated with Achilles Tendon Fatigue Damage in a Cadaveric Loading Model

**DOI:** 10.1101/849943

**Authors:** Elaine C. Schmidt, Todd J. Hullfish, Kathryn M. O’Connor, Michael W. Hast, Josh R. Baxter

**Affiliations:** Biedermann Lab for Orthopaedic Research, University of Pennsylvania, Philadelphia, Pennsylvania, USA; Department of Orthopaedic Surgery, University of Pennsylvania, Philadelphia, Pennsylvania, USA

**Keywords:** Achilles tendon, Ultrasound, Echogenicity, Biomechanics, Tendon Disorders

## Abstract

Achilles tendon disorders are among the most difficult sports-related injuries to predict with current diagnostic tools. The purpose of this study was to identify a clinically useful marker for early tendon damage. We hypothesized that alterations in mean echogenicity are linked with changes *in vitro* tendon mechanics. To test our hypothesis, we harvested Achilles tendons from 10 fresh-frozen cadaveric feet and cyclically fatigued them using a universal test frame while we continuously acquired ultrasound images. Throughout this fatigue protocol, we applied 2 stress tests every 500 loading cycles to quantify changes in ultrasound imaging echogenicity. We continued this fatigue protocol until each tendon either failed completely or survived 150,000 cycles. Tendons that failed during the fatigue loading (6/10) underwent greater changes in mean echogenicity compared to tendons that did not fail (*P* = 0.031). These tendons that failed during fatigue loading demonstrated greater changes in mean echogenicity that surpassed 1.0%; whereas survivor tendons exhibited less than 0.5% changes in mean echogenicity. We found that changes in mean echogenicity measured with ultrasound increased proportionally with increased tendon damage. The magnitude of these changes was relatively small (<1.5% change in mean echogenicity) but may be an effective predictor of tendon failure. Mean echogenicity is a promising marker for quantifying fatigue damage in cadaveric Achilles tendons during a stress test. Although these changes cannot be detected with the naked eye, computer-based predictive models may effectively assess risk of tendon damage in physically active adults.

**Level of evidence:** Controlled laboratory experiment

## INTRODUCTION

Achilles tendon disorders are among the most common conditions observed by sports medicine physicians and among the most difficult to predict with current clinical tools (McAuliffe et al., 2016). The incidence rates of acute and chronic Achilles injuries have increased proportionally with competitive and recreational sport participation over the past several decades (Järvinen et al., 2005). They are most commonly reported in athletic populations that experience either high-impact or repetitive lower extremity loading (Amin et al., 2013). Peak incidence occurs in individuals between 30 and 40 years of age (Lantto et al., 2015; Leppilahti et al., 1996). While Achilles tendinopathy is relatively manageable, one in twenty patients with tendinopathy will eventually sustain an Achilles tendon rupture (Yasui et al., 2017). This is likely caused by repetitive tendon strain and aggregation of microtears, which develop faster than they can be repaired (Järvinen et al., 2005). Despite improved treatment and rehabilitation protocols, these injuries can result in long-term deficits in patient function and mobility. Significant decreases in return to play, play time, and athletic performance have been extensively documented in elite basketball players (Amin et al., 2013; Deitch et al., 2006; Trofa et al., 2017). In a small percentage of cases these ruptures are career-ending - even following appropriate surgical care (Amin et al., 2013; Lorimer and Hume, 2014).

Tendon injuries are diagnosed based on clinical presentation and medical imaging results. While qualitative imaging is a validated method to grade the severity of tendinopathy (Sunding et al., 2016a), predicting the risk of these patients progressing or suffering tendon ruptures remains a major clinical need. Diagnosing chronic tendon injuries rely on qualitatively assessing tendon health with assays such as patient-reported measures of pain, fusiform swelling, and vascular infiltration of the tendon. However, more than half of complete Achilles tendon ruptures occur spontaneously and are not preceded by the same identifiable symptoms (Fredberg and Bolvig, 2002; Kannus and Józsa, 1991). Histological examination of these ruptures indicate that collagen degeneration, tenocyte necrosis, and acute inflammation are typically found near the rupture sites (Cetti et al., 2003). This suggests that many of the degenerative changes to the structure of the tendon have been initiated before a patient is able to seek treatment. To advance treatment and improve clinical outcomes, development of clinical imaging tools, which are sensitive enough to detect and quantify markers of pre-symptomatic tendinopathy and rupture are required (Järvinen et al., 2005; Kayser et al., 2005).

Ultrasound is currently used to diagnose tendon injury, and it is known that symptomatic tendon qualitatively provides less echogenicity - appearing darker on the image - than healthy tendon (Kainberger et al., 1990; Sunding et al., 2016b). However, to our knowledge, quantifiable changes in echogenicity have not been reported and the link between changes in echogenicity and fatigue loading in human tendons remains unknown. Therefore, the purpose of this study was to determine the efficacy of quantitative ultrasound imaging to explain *in vitro* fatigue-induced degradation of Achilles tendon mechanical properties. We hypothesized that decreases in mean echogenicity would be linked to *in vitro* tendon fatigue characterized by decreased mechanical properties. To test this hypothesis, we quantified the mean echogenicity and mechanical properties of cadaveric tendons during clinically relevant tendon stress tests throughout the fatigue life of tendons using a controlled cyclic loading protocol.

## METHODS

In this cadaveric tendon study, we dissected and prepared 10 cadaveric Achilles tendons from fresh-frozen lower leg specimens from 7 donors (Sex: 4M, 3F; Age: 60±15 years). We stored these specimens in a −20°C freezer and thawed overnight prior to dissection. For each specimen, we dissected out the entire Achilles tendon and its insertion into the calcaneus by cutting the bone in the sagittal plane at the calcaneal neck. Excised tendons were kept hydrated in saline-soaked gauze and re-frozen, which does not affect tissue properties (Jung et al., 2011). On the day of testing, we cut 3 cm long dog-bone shapes into the still-frozen tissue approximately 2-4 cm proximal to the calcaneus insertion (Figure 1A). To ensure these dog-bone cuts were reproducible, we used a #10 scalpel and a custom 3D-printed template as a guide. Although this process changed the native geometry of the tissue, it ensured that specimens would fail at the mid-substance (Woo et al., 1983) where our ultrasound images were acquired. To improve grip strength during testing, we embedded the calcaneal fragments in 45 × 35 × 55 mm blocks of bone cement (Poly(methyl methacrylate) (PMMA)). A transverse hole was drilled through the cement encasing the calcaneal bone for fixturing purposes.

**Figure 1.**
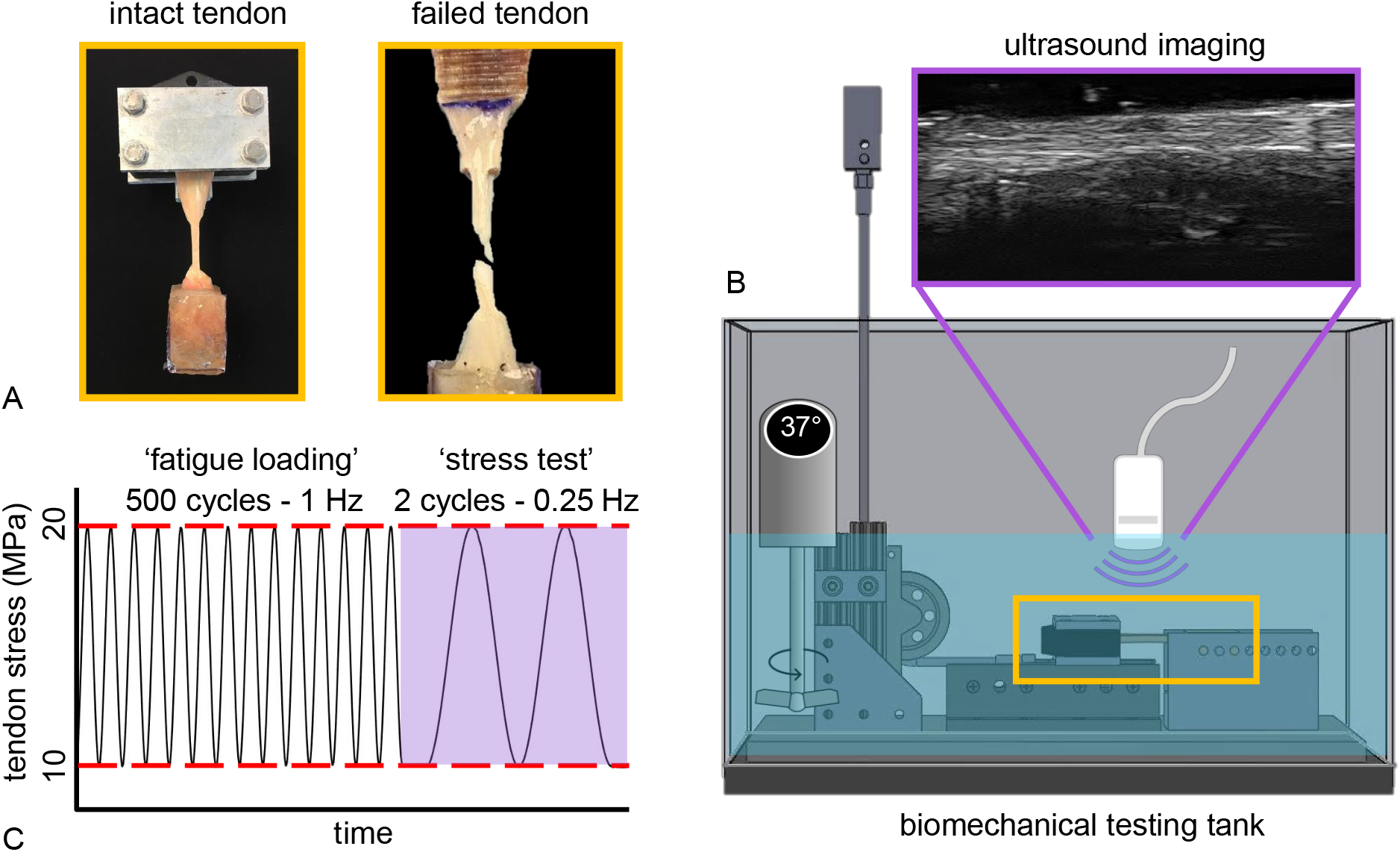
**(2 column)**: (A) We cut a dog-bone shape into each Achilles tendon specimen to ensure that tendon damage would occur in the mid substance. (B) These prepared specimens were then cyclically loaded in a custom-built tank and submerged in a warm solute bath to acquire ultrasound images throughout fatigue life. (C) To mechanically fatigue the test specimen and acquire clinically-relevant images, we applied 500 fatigue loading cycles at 1Hz followed by 2 stress test cycles at 0.25 Hz and repeated this loading protocol until the specimen either underwent 150,000 fatigue loading cycles or suffered complete mechanical failure.

We secured each test specimen to a custom-built tank and pulley system mounted on a universal test frame (Electroforce 3330, TA Instruments). The proximal end of the tendon was secured in custom-3D printed clamps reinforced with carbon-fiber filaments. Our previous experience dynamically actuating the Achilles tendon in cadaveric settings have shown that this type of clamping can withstand well above 1000 N (Baxter et al., 2016). This clamp was integrated into a linear track, which was anchored to the base of the tank to prevent rotation of the tendon during testing (Figure 1B). To secure the distal end of the specimen, we passed a stainless steel rod through potted calcaneal fragment and an aluminum carriage, which we anchored to the base of the tank. We connected the test specimen to the actuator frame using a braided wire-rope that we routed through a pulley system. Once secured, we submerged tendons in a temperature controlled (37°C) buffer solution (Phosphate-buffered saline) bath.

Prior to fatigue testing, we measured the cross-sectional area of each specimen using an 18 MHz B-mode ultrasound probe (L18-10L30H-4, SmartUS, Vilnius, Lithuania) and imaging software (EchoWave II 3.7.2 RC2). We measured this at the center of the dog-boned section under nominal tension (≈50 N), and used this measurement to transform a consistent stress profile to tendon-specific forces. While tendons were held under nominal tension (≈50 N), we positioned the ultrasound probe over the mid-substance of the dog-boned tendon (Figure 1B). The probe was held it in place with a custom 3-D printed mold and a chemistry clamp. Fine adjustments to the probe placement were made to ensure tendon fascicles were in line with the imaging plane and that the middle of the dog-boned portion was centered in the ultrasound image.

To fatigue each test specimen, we applied loading cycles that were generated using a sine waveform that cycled between 10-20 MPa at 1 Hz (Figure 1C). Using a digital trigger and custom script (MATLAB 2017b, The Mathworks Inc, Natick, MA), the ultrasound probe automatically acquired continuous images of the mid-substance of the dog-boned tendon after the completion of every 500 cycles. During this window of image acquisition, 2 loading cycles were performed at 0.25 Hz to eliminate motion artifact, during which we continuously acquired ultrasound images at 41 Hz. This process was repeated for a total of 150,000 cycles or until mid-substance failure occurred. Mechanical data were logged at 20 Hz for the entirety of the fatigue testing protocol. These data were used to calculate the absolute change in strain during the 0.25 Hz loading cycles across the fatigue life.

To analyze the series of ultrasound images, we used a custom image analysis algorithm to determine the mean echogenicity of each specimen. First, we assessed images at 10 MPa minima of stress throughout the fatigue life of the tendon. Second, we quantified changes in echogenicity during periods of elongation caused by increasing tendon stress from 10 MPa to 20 MPa during the 0.25 Hz stress tests. This measure of change in echogenicity during a slowly applied load is clinically analogous to a stress test. We synchronized each ultrasound video of the mid-substance with the mechanical testing data using the digital trigger and then the first frame of each video (where the tendon was dwelling at 10 MPa). To ensure consistent tendon analysis, one of our investigators manually identified rectangular regions of interest for each test specimen. Prior to analysis, we defined this region of interest to include as much of the tendon as possible, making sure to omit the deep and superficial edges of the tendon, which are highly echogenic. We then analyzed each ultrasound image using established methods to calculate echogenicity of Achilles tendon (Hullfish et al., 2018). Briefly, greyscale ultrasound images are made up of pixels with values between 0 and 255, with 0 (black) representing a hypoechoic medium such as water, and 255 (white) representing hyperechoic medium such as bone. Mean echogenicity of the tendon was calculated as the average greyscale value of all of the pixels within the region of interest in each video frame. Because we acquired all images using the same scan parameters, we divided the raw mean echogenicity values by 256 and reported as the percentage of maximum echogenicity. Additionally, the difference in mean echogenicity between each frame was calculated to determine how changes in mean echogenicity were related to changes in strain during the 0.25 Hz loading cycles. To understand how this relationship changed across fatigue life, we calculated the change in mean echogenicity as the difference between mean echogenicity at 10 and 20 MPa for each cycle (Figure 2A) over the course of fatigue life (Figure 2B).

**Figure 2.**
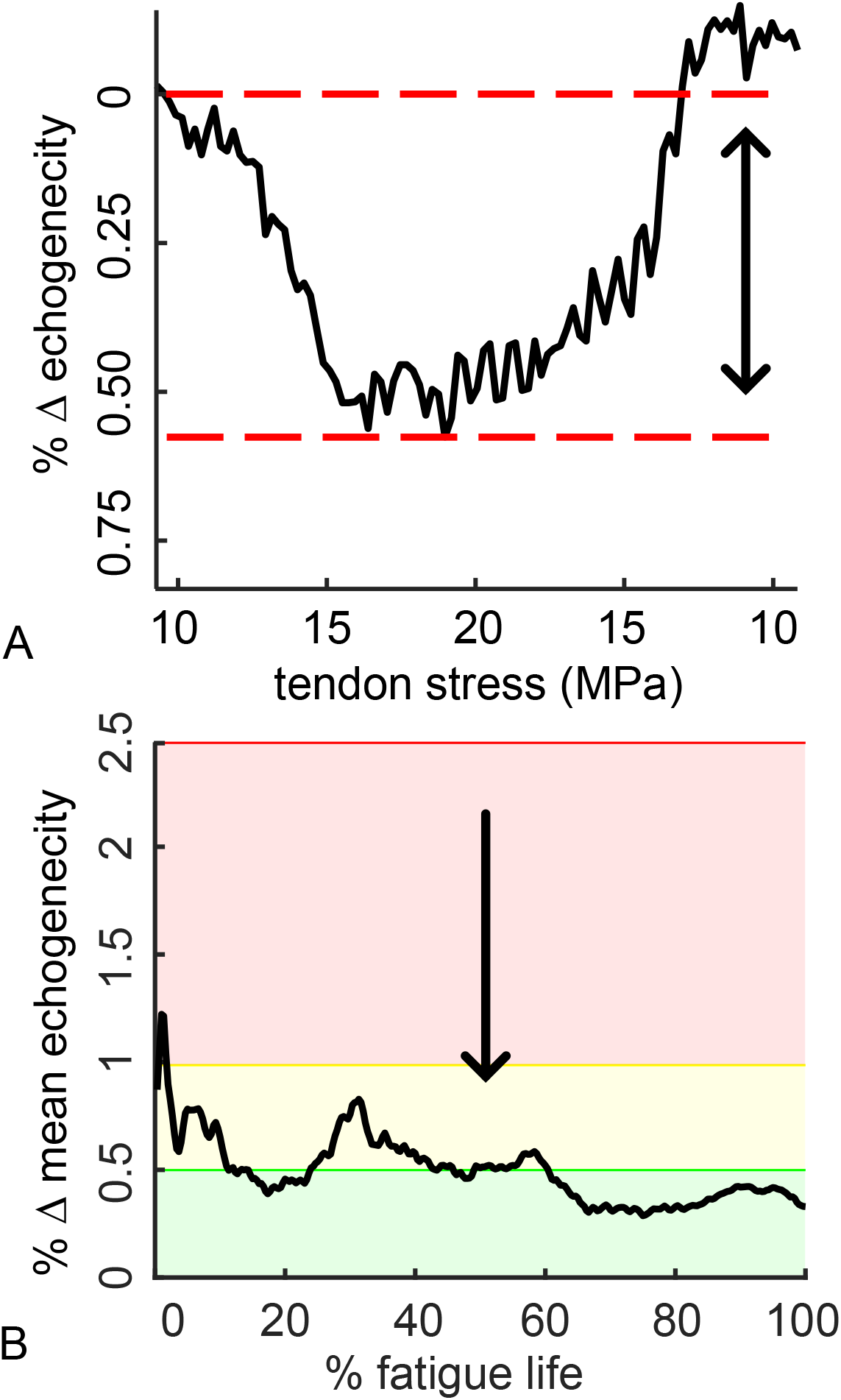
**(1 column)**. (A) During each stress test, we calculated the percent change in mean echogenicity (ΔME). (B) Plotting these changes throughout the fatigue life of the specimen highlights the structural changes we quantified using ultrasound imaging. To show the structural fatigue zones, we highlighted that less than 0.5% ΔME as green, between 0.5% and 1% ΔME as yellow, and greater than 1% ΔME as red.

To determine if changes in mean echogenicity and change in strain differed between tendons that failed and tendons that survived the fatigue loading, we compared the two groups using an unpaired t-test (p < 0.05). This comparison was made using the average change in mean echogenicity and average change in strain across the second phase of fatigue life to avoid any errors introduced by the pre-conditioning of the tendon in the first phase of fatigue life. Based on literature reports that found sex does not affect Achilles tendon properties (Morrison et al., 2015), we did not consider sex as a statistical variable.

## RESULTS

Of the ten tendons that were tested, six failed at the mid-substance and four did not fail after 150,000 cycles (Figure 3). Tendons that did fail were older on average (65 ± 11 years) than the tendons that did not fail (53 ± 13 years). The number of cyclic loads that caused tendon failure was highly variable (44,660 ± 22,269 cycles). Average normalized strain was 12.5±1.1% for tendons at failure and 7.1±1.6% for tendons that did fail. Failure occurrence appeared to be tendon-specific and independent of the donor, as both tendons from one matched pair (1L/R) failed while the contralateral tendons from (2L, 3L) did not.

**Figure 3.**
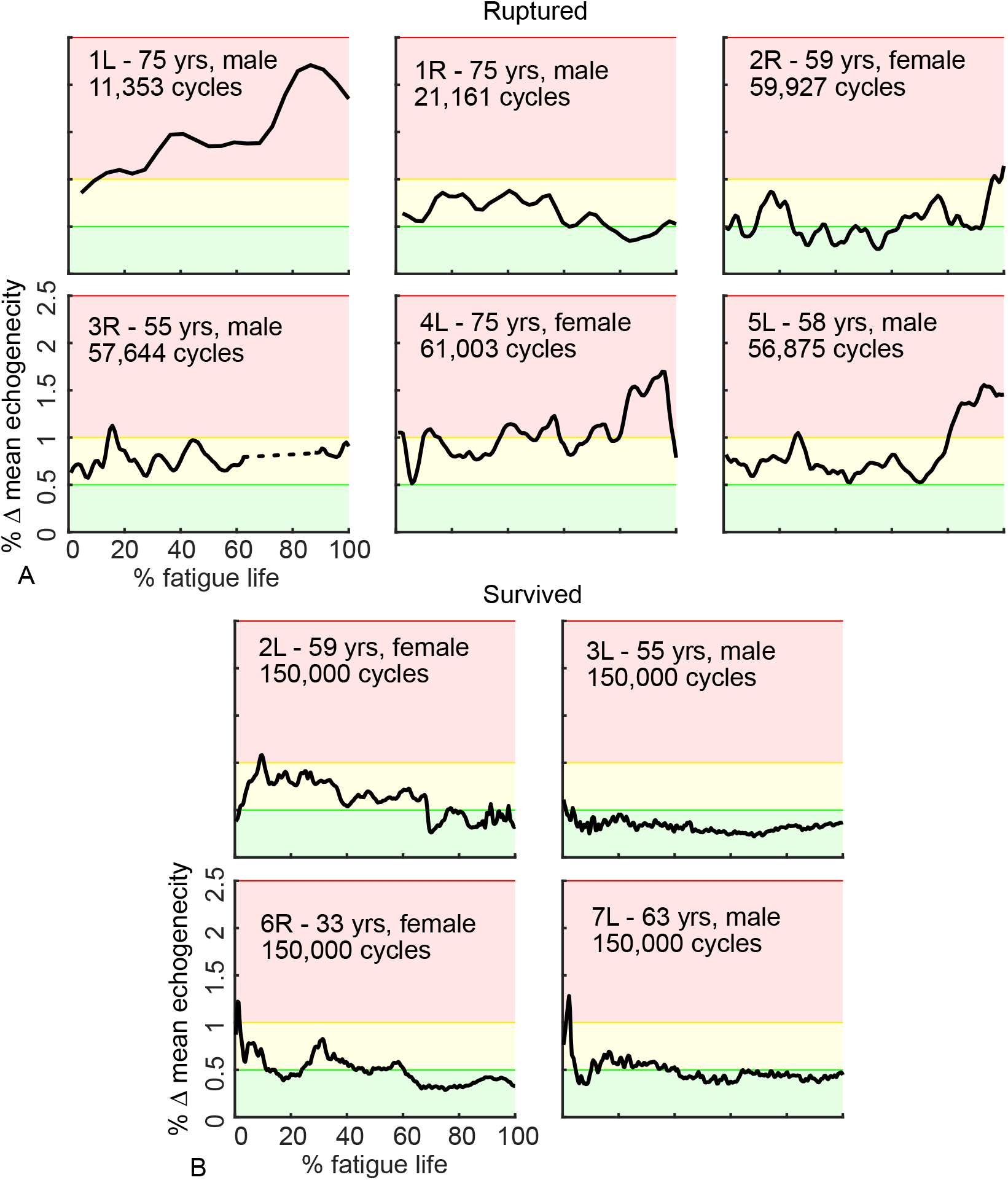
**(1.5 column)**. Changes in mean echogenicity (ΔME) during each 0.25 Hz waveform differed between (A) tendons that ruptured and (B) tendons that survived the 150,000 cycles. The three shaded zones are the result of a post-hoc examination of change in mean echogenicity. For specimen 3R, the dashed line indicates where ultrasound imaging did not occur due to an unforeseen complication with data collection.

Quantitative analysis of the ultrasound images indicated 2 key differences between tendons that failed during the cyclic loading protocol and those that did not. First, mean echogenicity decreased before failure. Second, the average change in mean echogenicity was significantly greater in tendons that failed (Figure 4A, *P* = 0.031). For most tendons that did fail, mean echogenicity decreased during the third phase of fatigue life. For the tendons that did not fail, mean echogenicity plateaued along with strain during the second phase of fatigue. Similarly, changes in mean echogenicity plateaued in tendons that did not fail while tendons that did fail demonstrated larger changes in mean echogenicity. Elevated changes in mean echogenicity throughout fatigue life (>1.0%) was associated with all tendons that failed the fatigue loading. Conversely, the tendons that survived the fatigue loading exhibited changes in mean echogenicity below 0.5% (Figure 3). In general, failing tendons exhibited changes in mean echogenicity of 0.5% or greater across the majority of fatigue life. These tendons that ruptured also underwent greater strain throughout their fatigue life compared to the tendons that survived (Figure 4B, *P* = 0.038).

**Figure 4.**
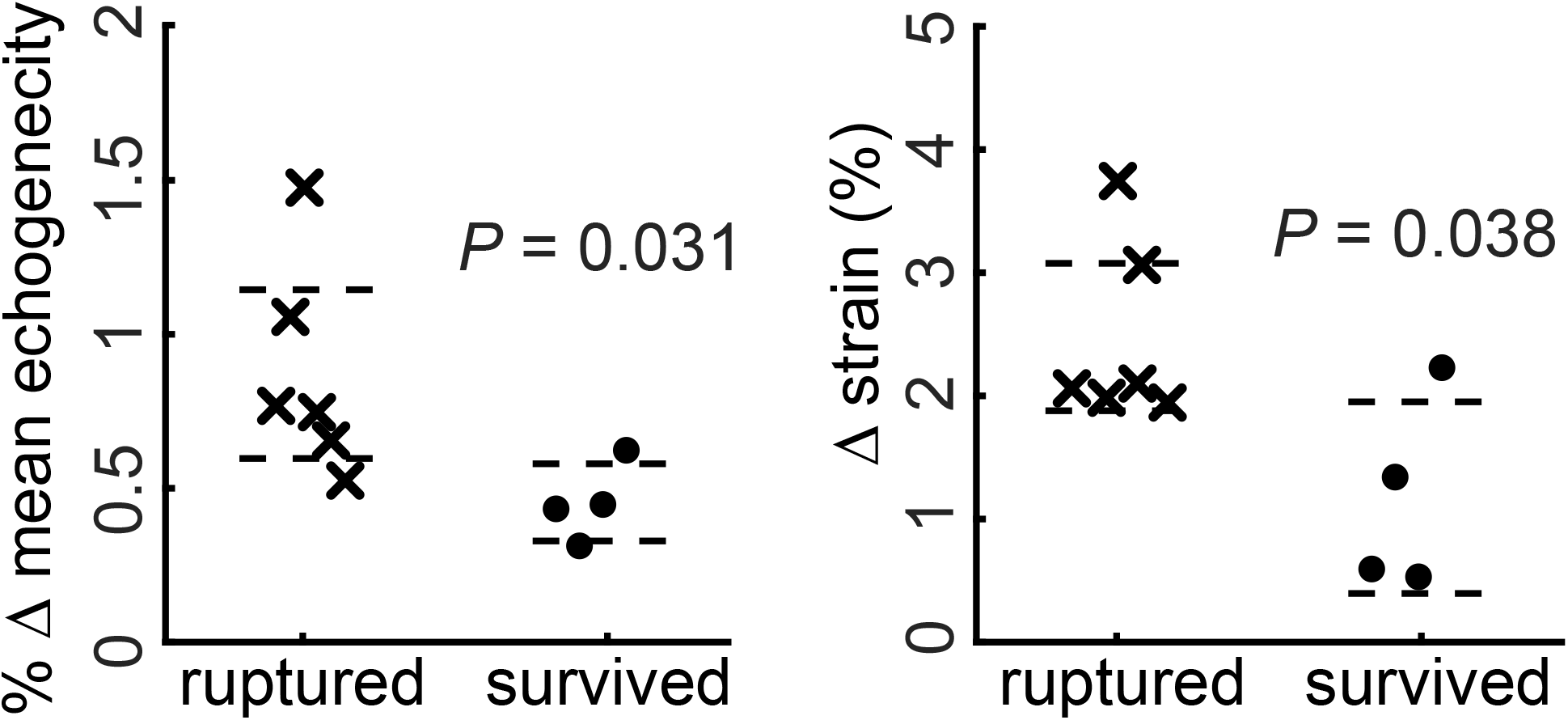
**(1 column)**. Differences for average change in (A) mean echogenicity and (B) average change in strain between the failure (crosses) and non-failure (filled circles) groups. Dashed lines indicate 95% confidence intervals.

## DISCUSSION

This study developed a novel model that tested cadaveric tissue with a controlled biomechanical protocol, while assessing changes in ultrasound signals. We mechanically fatigued cadaveric Achilles tendons and found that changes in mean echogenicity during a stress test appear to serve as reliable indicators of increased tendon damage. Similar studies on the material properties of Achilles tendons have tested whole cadaveric tendons at room temperature (Wren et al., 2001) or have used ultrasound imaging in human subjects synchronized to dynamometry (Arya and Kulig, 2010) or motion capture (Lichtwark and Wilson, 2005). To the best of our knowledge, the only other studies that have examined *in vitro* Achilles tendon damage with ultrasound synchronized to load data have been conducted in small animal models with a focus on repair of tendon injury (Freedman et al., 2014; Riggin et al., 2014). The present study represents the first attempt to quantify mean echogenicity in human cadaveric tissue as a marker of tendon damage. Compared to other methods of assessing tendon health, ultrasound is an attractive clinical tool because it is relatively inexpensive, portable, and non-invasive. Additionally, it is already used by clinicians to diagnose tendinopathy and tendinosis. The effective use of quantitative ultrasound to identify patients and athletes that are at risk of Achilles tendon rupture before an injury would be a major advancement.

Machine learning represents one future direction for our research, which aims to establish a predictive model for Achilles tendon injuries. Machine learning has distinct advantages, which could be directly applied to the results of this study. For example, the magnitude of change in mean echogenicity found in this study was very small (1%) and is not visible to the clinician’s eye. In our highly controlled experiment, eliminated the need for automated detection algorithms. However, these approaches approach may become necessary to quantify tendon health and fatigue damage in clinical settings. Within the past decade, machine learning has been impactful on the predictive power of medical imaging in many different areas of medicine (Greenspan et al., 2016; Wernick et al., 2010). Our future work is aimed at detecting small changes in mean echogenicity during tendon stress tests to identify patients at risk of further tendon injuries. Attempts to predict human Achilles tendinopathy *in vivo* have already started to yield some promising outcomes. A recent study used degree of speckle patterning in ultrasound imaging to classify Achilles tendon images as tendinopathic or healthy and demonstrated a prediction accuracy of greater than 80% (Bashford et al., 2008). Change in mean echogenicity may be a useful metric to add to these predictive models and could provide more accurate classification and quantification of the degree of tendon disorganization.

This study had several limitations that should be considered. Due to the cadaveric nature of this experiment, no biological responses to repetitive injury, such as inflammation or healing, could occur. This could affect interpretation of long-term fatigue on the tendons, and thus, results for total cycles to failure should be interpreted with some reservation. However, prolonged tendon loading within a very short period of time – for example, during distance running – likely mechanically stresses the tendon without stimulating an immediate biologic response. The tendons in this study were loaded in a purely horizontal direction from calcaneal insertion to clamp interface. This does not completely recapitulate the same motions between collagen fascicles that arise when the Achilles tendon is placed in an angle closer to the anatomical range or under muscle force imbalances that could originate from the triceps surae (Lersch et al., 2012). There are three distinct bundles of collagen fascicles that arise from the medial and lateral gastrocnemius and the soleus that each twist helically by varying amounts as they insert distally into the calcaneus. This may cause the non-uniform motions and relative sliding between fascicle bundles that have been observed through ultrasound analysis of the Achilles under passive and eccentric loading *in vivo* (Slane and Thelen, 2014). Although the dog-bone method for preparing the tendons was utilized to control failure precisely at the mid-substance of the tendons, where the ultrasound probe was focused, this may have also disrupted the natural sliding and twisting movements of collagen fascicle bundles. However, the strain measurements reported here are similar to a previous report that tested intact Achilles tendons (Wren et al., 2001). We did not combine our B-mode ultrasound imaging analysis with other assays for tendon damage that have been utilized in other studies on Achilles tendon mechanical or substructural properties, such as sonoelastography (Klauser et al., 2013), polarized light microscopy (Kainberger et al., 1990), analyzing collagen organization (Hullfish et al., 2018), or histology (Maffulli et al., 2000). This limits us from understanding what is happening dynamically at the microscopic level and prevents full characterization of the effects of heterogeneous distribution of tendon strain on failure. Due to limited sample size, we were not able to determine if donor age or gender affected total number of cycles to failure. However, our study was strengthened by variation in mechanical response to fatigue loading, as it highlighted the association between ultrasound imaging and tissue damage.

## CONCLUSION

Ultrasound imaging provides clinicians with an attractive opportunity to visualize structural changes that occur within a damaged tendon, but these changes are difficult to detect qualitatively before they become symptomatic. This study found detectable differences in image echogenicity during a stress test between tendons that fail during cyclic loading and those that do not. While preliminary, our findings indicate that B-mode ultrasound has potential as a clinically viable tool to predict severe tendon injuries. Our future work is focused on developing computer-based predictive tools to assess Achilles tendon fatigue in patients with tendinopathy following prolonged tendon loading to establish quantitative imaging thresholds that can serve as clinical benchmarks.

